# “Acute Respiratory Distress and Cytokine Storm in Aged, SARS-CoV-2 Infected African Green Monkeys, but not in Rhesus Macaques”

**DOI:** 10.1101/2020.06.18.157933

**Authors:** Robert V. Blair, Monica Vaccari, Lara A. Doyle-Meyers, Chad J Roy, Kasi Russell-Lodrigue, Marissa Fahlberg, Chris J. Monjure, Brandon Beddingfield, Kenneth S. Plante, Jessica A. Plante, Scott C. Weaver, Xuebin Qin, Cecily C. Midkiff, Gabrielle Lehmicke, Nadia Golden, Breanna Threeton, Toni Penney, Carolina Allers, Mary B Barnes, Melissa Pattison, Prasun K Datta, Nicholas J Maness, Angela Birnbaum, Tracy Fischer, Rudolf P. Bohm, Jay Rappaport

## Abstract

SARS-CoV-2 induces a wide range of disease severity ranging from asymptomatic infection, to a life-threating illness, particularly in the elderly and persons with comorbid conditions. Among those persons with serious COVID-19 disease, acute respiratory distress syndrome (ARDS) is a common and often fatal presentation. Animal models of SARS-CoV-2 infection that manifest severe disease are needed to investigate the pathogenesis of COVID-19 induced ARDS and evaluate therapeutic strategies. Here we report ARDS in two aged African green monkeys (AGMs) infected with SARS-CoV-2 that demonstrated pathological lesions and disease similar to severe COVID-19 in humans. We also report a comparatively mild COVID-19 phenotype characterized by minor clinical, radiographic and histopathologic changes in the two surviving, aged AGMs and four rhesus macaques (RMs) infected with SARS-CoV-2. We found dramatic increases in circulating cytokines in three of four infected, aged AGMs but not in infected RMs. All of the AGMs showed increased levels of plasma IL-6 compared to baseline, a predictive marker and presumptive therapeutic target in humans infected with SARS-CoV-2 infection. Together, our results show that both RM and AGM are capable of modeling SARS-CoV-2 infection and suggest that aged AGMs may be useful for modeling severe disease manifestations including ARDS.

## Introduction

The coronavirus disease-2019 (COVID-19) pandemic, caused by the novel coronavirus, severe acute respiratory syndrome coronavirus-2 (SARS-CoV-2), has resulted in the deaths of hundreds of thousands of people and has caused massive economic and health disruptions across the globe. This unprecedented level of disruption has been driven by two main features of SARS-CoV-2: its high rate of person-to-person transmissibility and the potential to cause severe, life-threatening, pneumonia. Although severe disease is only seen in a small subset of infected people (1), it is this outcome and minimal understanding of its pathogenesis that has resulted in global unrest. Research into the causes and mechanisms of the most severe manifestations of COVID-19 is needed to inform and facilitate the development of prophylactic and therapeutic approaches that can prevent this life-threatening outcome.

Infection with SARS-CoV-2 and development of COVID-19 is accompanied by a mild respiratory disease for most individuals. However, a small subset progress to develop severe respiratory disease which, in some cases, is fatal (1). The most severely affected individuals often present with a fever, cough, dyspnea, and bilateral radiographic opacities that, in the majority of critically ill patients, progresses to acute respiratory distress syndrome (ARDS) (2). The onset of ARDS is often associated with an increase in circulating pro-inflammatory cytokines (3,4). Interleukin-6 (IL-6), in particular, has been shown to correlate with radiographic scores in patients with SARS-CoV-2 infection (5). Worsening of disease can be seen in the context of declining viral loads and markedly elevated cytokines suggesting a role for these inflammatory responses in disease progression and immunopathology (6). Despite extensive research during both the SARS-CoV and the Middle East respiratory syndrome (MERS) outbreaks, the factors that drive this inflammatory response are still poorly understood.

Animal models have been used extensively during previous outbreaks of SARS-CoV (7–11) and MERS (12–14) to model disease progression and to test vaccines and therapeutics. Nonhuman primates (NHPs) are ideally suited to model respiratory human viral infections primarily because of the similarities to human respiratory anatomy and immunologic responses when compared to other animal species. Several NHP species have already been successfully employed to model pathogenesis (15–19) and test vaccine candidates (20–23) for SARS-CoV-2. These prior studies have shown NHPs are susceptible to infection and develop mild to moderate disease, but none has been able to recapitulate the rapid clinical deterioration seen in people with severe disease and ARDS. NHP models capable of recapitulating the entire spectrum of SARS-CoV-2 manifestations, from mild to severe disease, are urgently needed to not only test the efficacy of vaccines and medical countermeasures that are currently being developed in response to COVID-19, but to also investigate the pathogenesis and virus-host interactions of SARS-CoV-2. Age is a well-established risk factor for severe disease and death in humans infected with SARS-CoV-2 (2,24,25), and therefore we challenged older RM and AGMs with SARS-CoV-2 to see if a similar more severe disease phenotype was observed in aged cohorts.

Here we report the sudden and rapid health deterioration of two out of four aged AGMs experimentally infected with SARS-CoV-2. The two affected animals developed pneumonia, ARDS and increased plasma cytokines similar to the complications reported in 5-13% of COVID-19 patients (26).

## Results

### SARS-CoV-2 infection and viral kinetics in RM and AGM

Four, aged, AGMs and four RM, thirteen to fifteen years of age, were exposed by two routes to SARS-CoV-2 isolate USA-WA1/2020. Four animals were exposed via small particle aerosol (AGM1, AGM4, RM3, RM4) and received an inhaled dose of approximately 2×10^3^ TCID_50_. Four animals were exposed via multiple route installation (AGM 2, AGM3, RM1, RM2) including conjunctival, intratracheal, oral, and intranasal exposure resulting in a cumulative dose of 3.61×10^6^ PFU (Supplemental Table 1). SARS-CoV-2 RNA was detectable in swabs obtained from mucosal sites in all eight animals. For AGMs the viral RNA peaked between 3- and 7 days post inoculation (DPI) and persisted throughout the course of the study in pharyngeal and nasal swabs as well as bronchial brush samples (Figure 1). In RMs, viral RNA peaked earlier between 1- and 5 DPI. After peak, viral RNA loads in RM gradually declined to undetectable levels at all sites by 21 DPI, except in nasal swabs which had detectable virus at necropsy in two out of four RM. The highest levels of viral RNA were detected in the pharynx and nasal cavity with peaks at 10^7^-10^11^ and 10^8^-10^9^ in AGM and 10^6^-10^8^ and 10^5^-10^11^ copies per swab in RM, respectively. Rectal swabs contained high viral RNA loads similar to reports in humans (27,28); however, with dissimilar kinetics in AGMs relative to virus detected in other sites, peaking between 7- and 14 DPI. Viral RNA was also detected in vaginal swabs of the two female AGMs in contrast to reports in human subjects (29). Despite the 3 log difference in exposure dose, in comparing aerosol and mulit-route viral challenge, no significant difference was observed in the viral RNA loads or kinetics.

**Figure 1.**
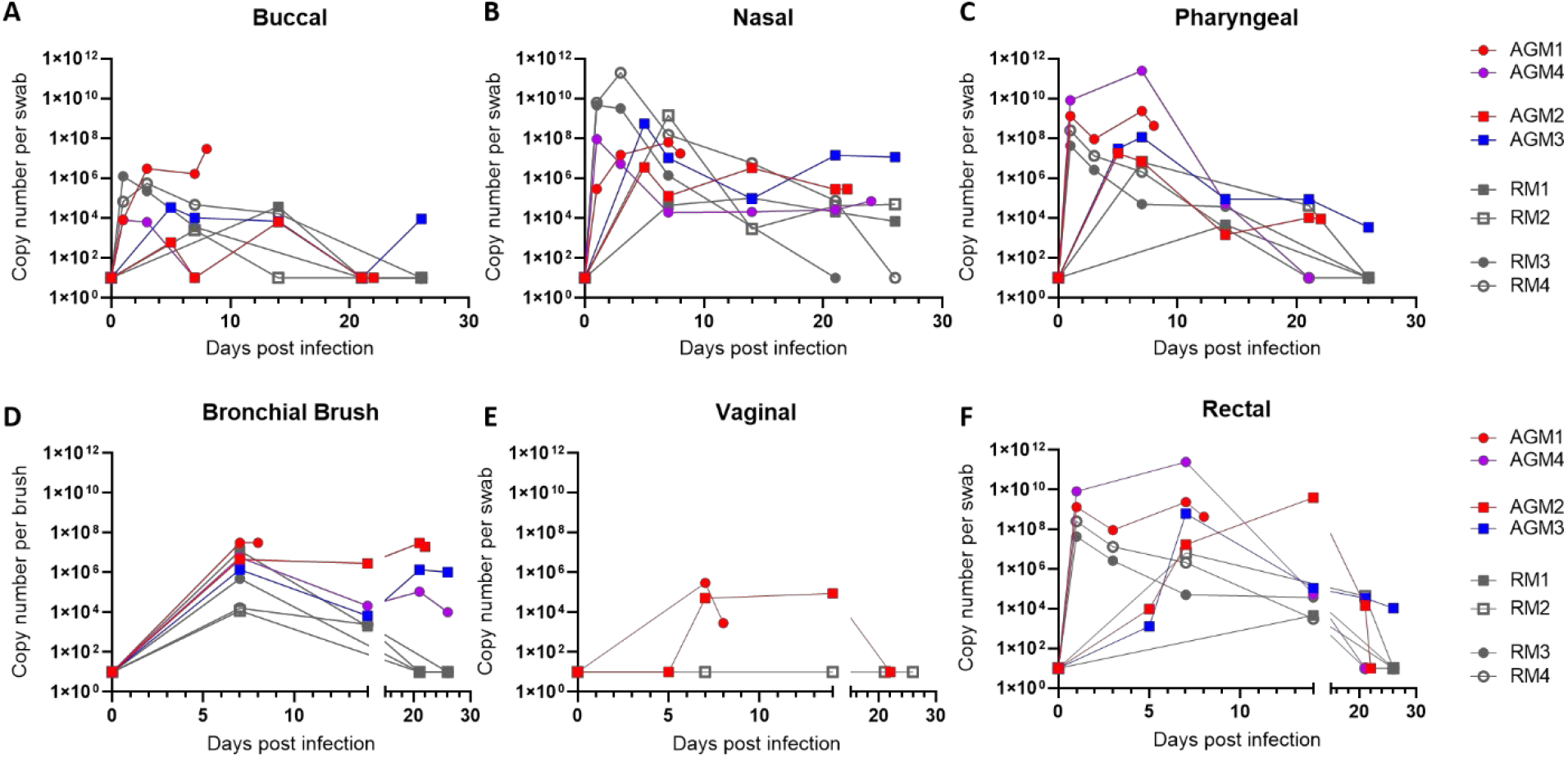
Quantification of viral RNA loads from mucosal swabs. All animals (4 African green monkeys and 4 rhesus macaques) had detectable virus at mucosal sites. No significant differences were noted in viral load between species and route of exposure (Mann-Whitney U test). Animals with ARDS trended to high viral loads in bronchial brush samples. Circles: aerosol exposure; Squares: multiroute exposure; Gray: Rhesus macaques; Color: AGM by outcome. Red: developed ARDS; Purple: increased cytokines without ARDS; Blue: no cytokine increase or ARDS.

### ARDS in two SARS-CoV-2 infected aged AGMs

After SARS-CoV-2 exposure, animals were followed up to four weeks post-infection with regular clinical assessment that included physical exam, pulse oximetry, and plethysmography. Clinical findings during the first 6 DPI included mild transient changes in SpO2 and appetite with one RM3 exhibiting mild intermittent fever (Supplemental Figure 1). At 7 DPI all animals underwent a complete physical evaluation and an extensive sample collection protocol including fluid (urine, CSF, BAL, and vaginal and rectal weks), stool, swab (buccal, nasal, and pharyngeal), and bronchial brush collection, no remarkable findings were noted. That afternoon (7 DPI), AGM1 developed mild tachypnea that progressed to severe respiratory distress in less than 24 hours. On the morning of the 8^th^ DPI, the animal was discovered recumbent and exam findings included dyspnea, tachypnea, hypothermia, and an SpO2 of 77% under oxygen supplementation (Supplemental Figure 1). Between day 8- and 21 DPI, mild transient changes in SpO2 and appetite were noted in all remaining animals, RM3 continued to have mild intermittent fever, and RM1 developed an intermittent cough. On 21 DPI all remaining animals underwent another complete evaluation. During the morning exam on 22 DPI, AGM2 began exhibiting tachypnea that progressed to severe respiratory distress by that afternoon. The onset, clinical presentation, and rate of progression of disease in AGM2 was similar to AGM1 and included dyspnea, tachypnea, hypothermia, and a Sp02 of 77% on ambient air. After 22 DPI, no significant clinical findings were observed in any of the remaining animals.

Thoracic radiographs for AGM1 and AGM2 revealed a diffuse alveolar pattern throughout the right lung fields and a lobar sign in the caudal dorsal lung field. In AGM2 the left caudal lung lobe also contained a mild alveolar pattern. These findings were in stark contrast to the radiographs from the day before highlighting the rapid disease progression (Figure 2). The radiographic presentation in severe human COVID-19 is similar and characterized by bilateral, peripheral, ill-defined ground glass opacifications that more frequently involve the right lower lobe (30). No radiographic changes were noted in AGM3 and AGM4. RM1 developed mild to moderate radiographic opacities in the right caudal lung lobe by 11 DPI which gradually resolved over time. RM2 revealed a small area of increased opacity in the ventral lung field at 11 DPI which was not observed on subsequent radiographs.

**Figure 2.**
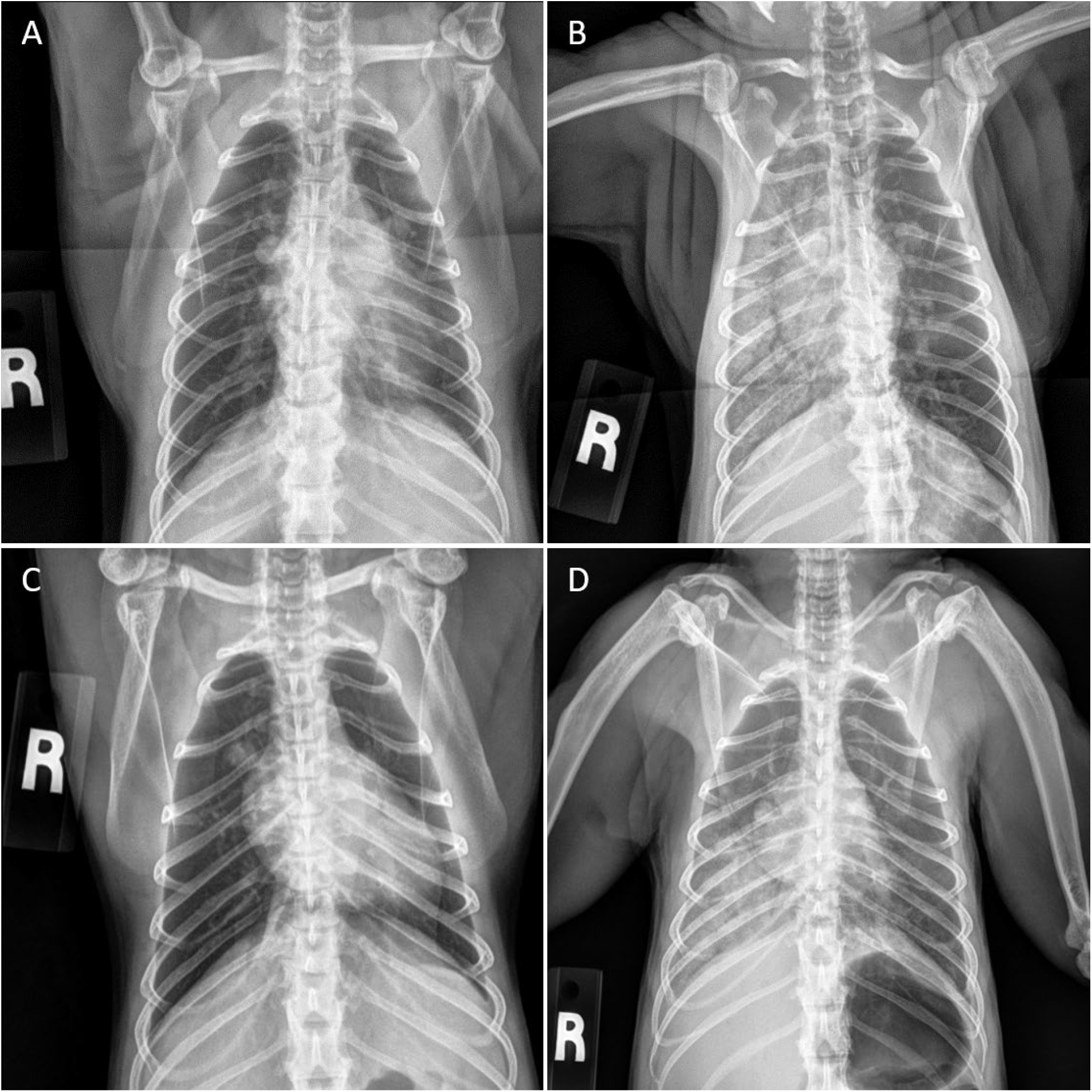
Radiographic changes in SARS-CoV-2 infected AGMs with ARDS. Radiographs the day prior (**A,C**) and at the time of necropsy (**B,D**) in AGM1 (**A,B**) and AGM2 (**C,D**) showing the rapid development of alveolar lung opacities within the lungs of both animals.

Rapid disease progression in AGM1 and AGM2 was associated with an elevated WBC, a mature neutrophilia and a normal lymphocyte count that resulted in an elevated neutrophil-to-lymphocyte ratio (NLR) (Supplemental Figure 2). These changes were not observed in the other two AGM or in four RM. Mild lymphopenia was observed at acute time points in RM1-3; however, in the absence of neutrophilia, the NLR was only mildly elevated (<4) in these animals. Serum chemistries revealed hypoproteinemia, an elevated glucose, and mild to moderate elevation in BUN for both animals with ARDS. Creatinine, and AST were also mildly elevated in AGM1 indicating multiple organ dysfunction (Supplemental Figure 2). These chemistry abnormalities were not observed in the two remaining AGMs or four RM. All four RM developed mild hypoalbuminemia after infection, and three of the four developed mild hyperglobulinemia. Elevated NLR has been identified as an independent risk factor for mortality in hospitalized patients with SARS-CoV-2, and increased NLR significantly correlated with elevations in AST, glucose, BUN, and creatinine in these patients (31). The constellation of hematologic changes in AGM1 and AGM2 is therefore similar to the changes observed in human COVID-19 patients with increased NLR; however in humans, increased NLR is often associated with lymphopenia (32) which was not observed in these two animals.

Due to their rapidly declining clinical condition, AGM1 and AGM2 were euthanized at 8 and 22 DPI, respectively. All remaining animals (2 AGM and 4 RM) were euthanized at the study endpoint between three- and four-weeks post infection. A complete necropsy was performed on all animals.

### Increased plasma cytokines in two AGMs with, and one without, ARDS

Increased plasma cytokines has been observed in a subgroup of patients with severe COVID-19 pneumonia (33). In these patients the disease progresses rapidly, and mortality is high. A panel of cytokines was measured in plasma at baseline and during the course of infection. Interferon gamma (IFNγ) responses increased at 1 week post infection in all of the AGMs and none of the RM, as shown by the heat map (Figure 3A). IFNγ levels were higher in AGM1 and AGM2 and were associated with viral RNA in the bronchial brushes at the same time point (1 week) (Figure 3B, C).

**Figure 3.**
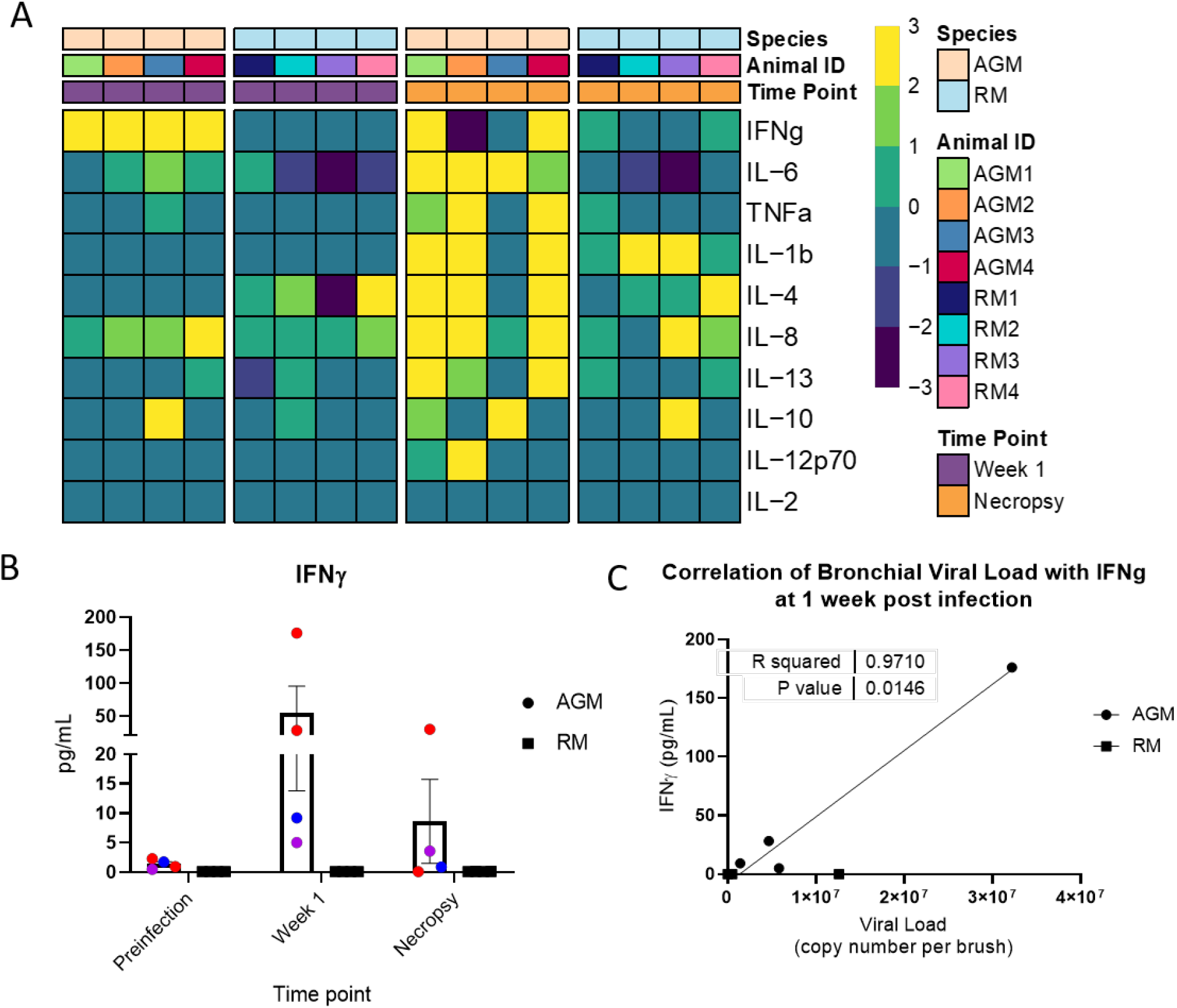
Cytokine increase. Heat map (**A**) showing changes in the levels of ten cytokines in plasma at week 1 and necropsy with respect to the baseline in AGMs and RM. Data are normalized (log2). **B**) Levels of IFNg (pg/ml) in plasma at baseline, week 1, and at necropsy. Column represents mean, error bars are SEM. **C**) Association between IFNg levels at week 1 and viral load in bronchiolar brushes (Pearson test).

A group of cytokines similar to those observed in human COVID-19 was upregulated in the two animals that progressed to ARDS (AGM1 and AGM2) at the time of necropsy compared to baseline levels (Figure 3A and Supplemental Figure 3). Elevated markers included IFNγ, IL-6, IL-4/IL-13, IL-8, IL-1β and TNFα. AGM4 did not develop ARDS; however, this animal showed a similar increase in cytokine concentrations, but with only a mild elevation of IL-6. AGM3 had increased levels of IL-10 both at week 1 and necropsy and was notable for having the least severe histopathologic changes of the AGMs. In contrast, RM only showed modest changes in cytokine expression at one week post infection and necropsy, despite comparable peak viral loads.

### Antibody titers in SARS-CoV-2 infected AGM and RM

Binding IgG antibody to S1/S2 subunits of the spike (S) protein and nucleoprotein (NP) were measured longitudinally for all animals by ELISA and Multiplexed Fluorometric ImmunoAssay (MFIA), respectively. Antibodies were not detected prior to infection in any of the animals used in this infection study. In AGM1 (euthanized 8 DPI) antibodies were not detected. In all other animals antibodies to S and NP were detectable by 14 DPI except RM3 who did not have detectable antibodies to S until 21 DPI. No significant differences in antibody responses were noted by route or species. (Supplemental Figure 4).

### Pulmonary pathology in AGMs with ARDS

Gross postmortem examination of AGM1 and AGM2 revealed severe consolidation and edema in the right caudal lung lobe with generalized failure to collapse of the remaining lobes, consistent with a bronchointerstitial pneumonia (Figure 4). In AGM2 (multiroute exposure) pulmonary hemorrhage was also noted near the dorsal margin of the right caudal lung lobe (not shown). AGM3 had multifocal pleural adhesions between the left caudal lung lobe and the diaphragm that was interpreted as being unrelated to SARS-CoV-2 infection based on the chronicity of the lesion and the history of the animal. RM1 had a focal pulmonary scar in the right caudal lung lobe surrounded by acute hemorrhage (Supplemental Figure 5). The lungs of the remaining animals (AGM4, RM2, RM3, and RM4) were grossly normal. No gross abnormalities were noted outside the lungs in any of the eight animals.

**Figure 4.**
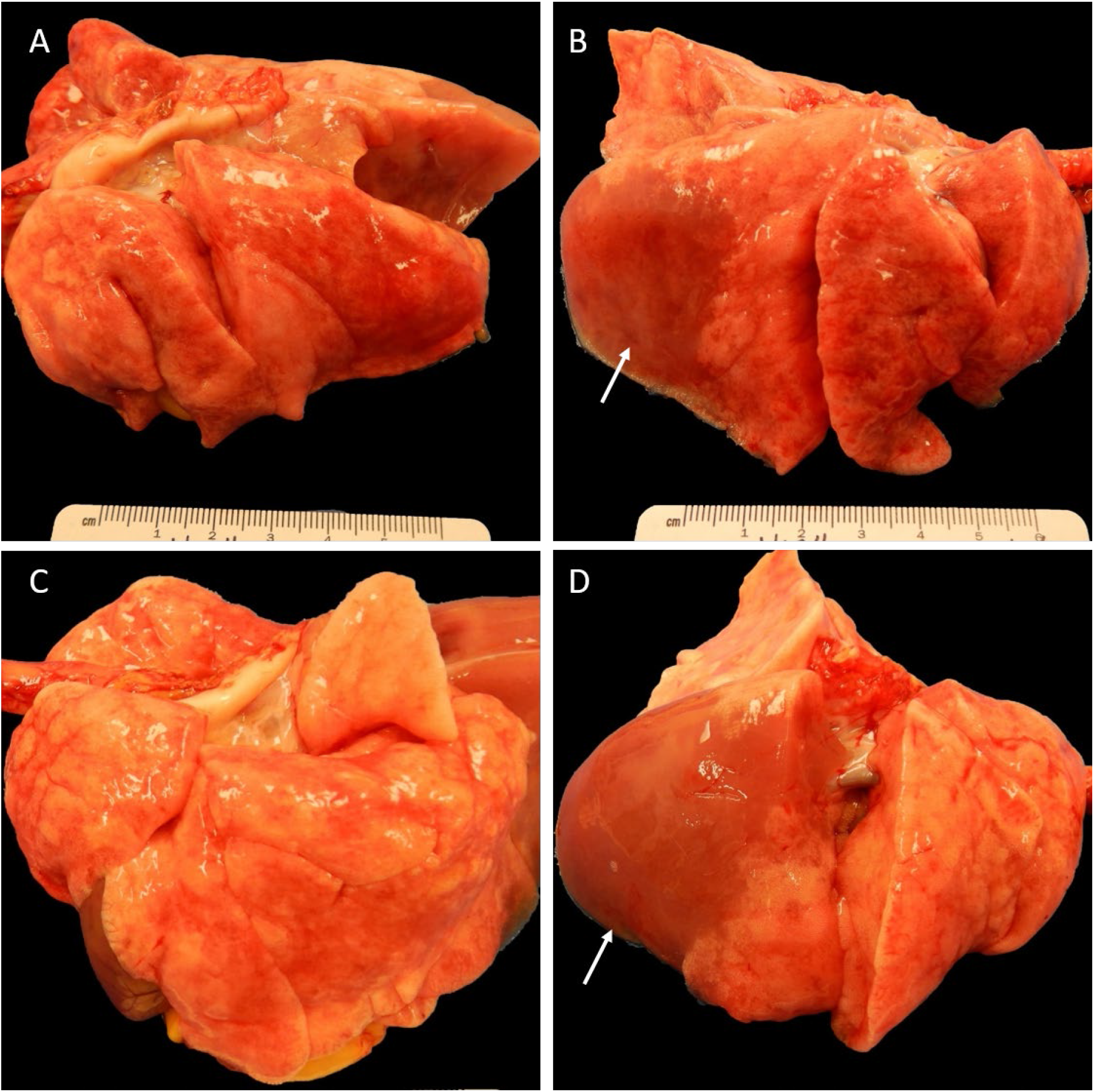
Gross pathologic findings of AGM with ARDS. Both AGM1 (**A,B**) and AGM2 (**C,D**) have extensive consolidation of the right caudal lung lobe (**B,D**; arrows) with lesser involvement of the right middle and cranial lung lobes. The left lung of AGM1 (**A**) fails to collapse. The left lung of AGM2 (**C**) does not have gross abnormalities.

Histopathologic findings in the lungs of AGM1 and AGM2 were similar and characterized by alveoli that were filled with fibrin, hemorrhage, and proteinaceous fluid. Alveoli were multifocally lined by hyaline membranes and/or type II pneumocytes, consistent with diffuse alveolar damage. Bronchial and alveolar septal necrosis were present within severely affected lung lobes characterized by a loss of epithelial lining cells and infiltration by neutrophils with lesser numbers of lymphocytes and histiocytes (Figure 5A, B). In AGM1, type II pneumocytes frequently exhibited atypia and occasionally contained mitotic figures. Regions of the lung from AGM1 also had organization of intra-alveolar fibrin with infiltration by spindle cells and lining by type II pneumocytes, consistent with early organizing pneumonia. Low numbers of multinucleated giant cell syncytia were scattered throughout alveoli (Figure 5C). Fluorescent immunohistochemistry identified low numbers of type II pneumocytes and alveolar macrophages that were positive for SARS-CoV-2 nucleoprotein within the affected lungs from AGM1, but not AGM2. (Figure 5D). AGM4 had multifocal, mild to moderate, interstitial pneumonia scattered throughout all lung lobes characterized by a mixed infiltrate of neutrophils, lymphocytes, and histiocytes. Multinucleated giant cell syncytia and atypical pneumocyte hyperplasia were rarely observed in all lung lobes. AGM3 had scant inflammation in all lung lobes examined (Supplemental Figure 6). Three out of four rhesus macaques (RM1, RM2, and RM4) had microscopic evidence of aspiration pneumonia characterized by a foreign plant material within bronchioles. Affected bronchioles were surrounded by mild (RM2 and RM4) or severe (RM1) pyogranulomatous inflammation (Supplemental Figure 7). In RM1, microscopic examination of the pulmonary scar noted grossly identified a markedly ectatic bronchiole surrounded by pyogranulomatous inflammation in the pulmonary parenchyma adjacent to the scar (not shown). This lesion was presumed to be secondary to aspiration, although no foreign material was identified within the tissue section. The right middle lung lobe of RM3 had moderate lymphocytic vasculitis with medial thickening of affected vessels. Viral load was not significantly associated with pulmonary pathology in AGMs or RMs; however, the two animals that developed diffuse alveolar damage showed higher viral loads in bronchi. Histopathologic lesions in other tissues were mild and interpreted as not significant in all eight animals (Supplemental Table 2).

**Figure 5.**
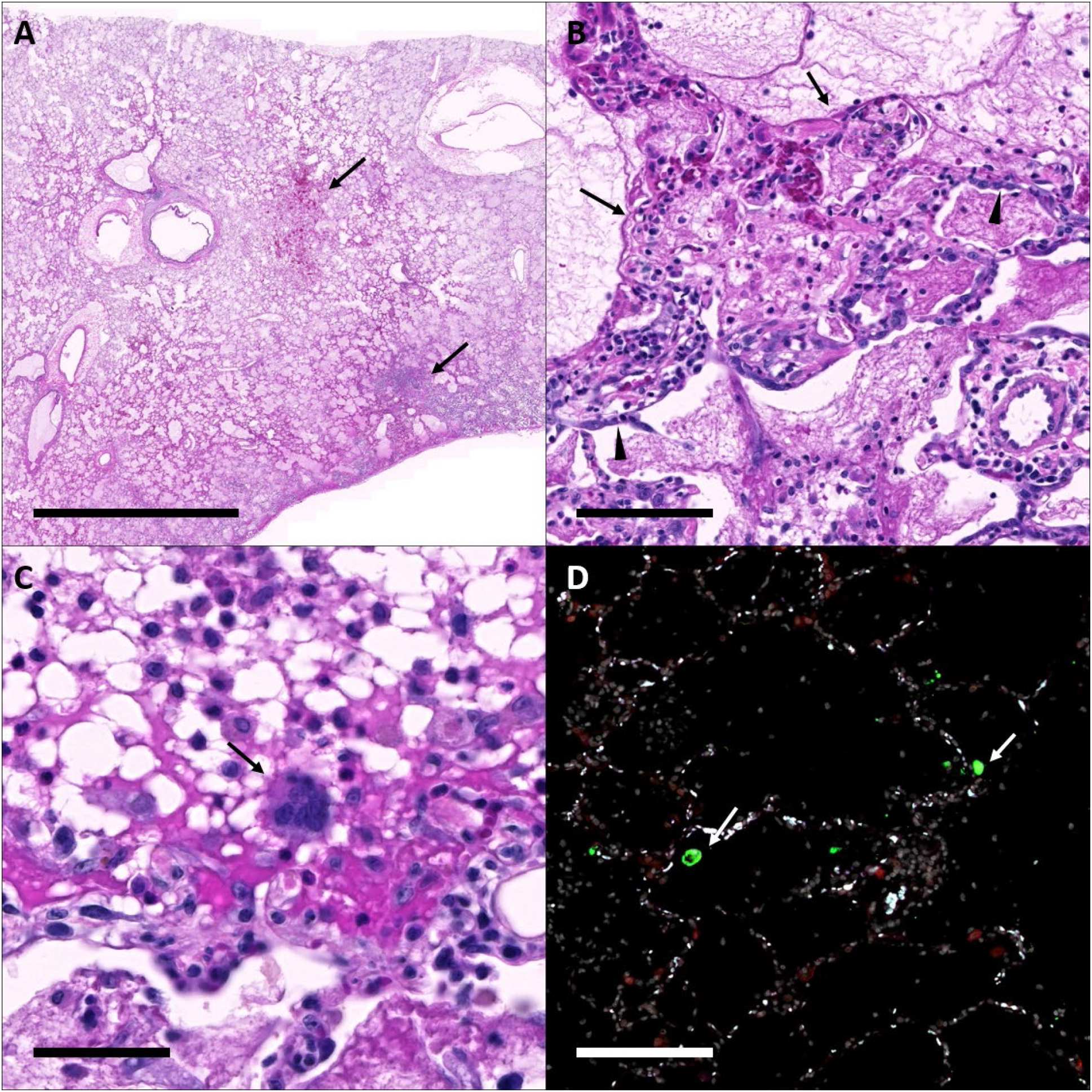
Histopathology and fluorescent immunohistochemistry in AGM1. (**A**) The right lower lung lobe is filled with fibrin and edema with areas of hemorrhage and necrosis (arrows); Bar = 5 mm. (**B**) Alveoli are variably lined by hyaline membranes (arrows) and type II pneumocytes (arrowheads); Bar = 100 um. (**C**) Rare multinucleated syncytia (arrow) are scattered throughout the affected lungs; Bar = 50 um. (**D**) Fluorescent immunohistochemistry for COV-2 nucleoprotein (green, arrows) and ACE2 (red) identified low numbers of CoV-2 positive cells within the affected lung lobes; Bar = 100 um. White: DAPI/nuclei; Green: CoV-2; Red: ACE2 Blue: Empty.

## Discussion

NHPs are ideal candidates for the modeling of human respiratory viral infections. Several recent studies have been published utilizing AGMs (15,18) and RMs (16,17) to model SARS-CoV-2 infection and have shown both species are capable of modeling mild to moderate disease and are useful for testing prospective vaccines and therapeutics. However, none of these prior studies have been able to recapitulate the spontaneous, severe disease phenotype seen in a subset of people with COVID-19. Our results show that following infection with SARS-CoV-2, aged AGMs can spontaneously develop ARDS and a cytokine release syndrome similar to that reported in humans with severe COVID-19 (33), which was not observed in RM of similar age.

Acute respiratory distress syndrome is defined by the rapid development (within 1 week of new respiratory symptoms) of bilateral radiographic opacities, and respiratory failure not explained by cardiac failure or fluid overload (34). Objective criteria that have been used for the diagnosis and scoring of ARDS include the number of lung quadrants affected on radiographs, the PaO2/FiO2 ratio, and measurements of positive end-expiratory pressure (PEEP) and pulmonary compliance (35). Two of the AGMs in our study developed widespread radiographic opacities and severe respiratory distress (Sp02 77%) that progressed over a 24-hour period. The rapid progression of clinical disease in conjunction with pre-established humane endpoints, precluded further diagnostics such as PEEP, pulmonary compliance, and echocardiogram; however postmortem examination found no evidence of congestive heart failure. Taken together the radiographic changes, impaired oxygenation, and the postmortem findings which included diffuse alveolar damage, are consistent with a diagnosis of ARDS in both AGM1 and AGM2. Of note, both of the animals that developed ARDS did so within 24 hours following routinely scheduled sampling procedures which included anesthesia and bronchoalveolar lavage. It has been our experience that these procedures are well tolerated, and procedure-related complications are exceedingly rare (fatal complication within 48-hours of procedure after 2 of 11,431 procedures in animals ranging from 1-31 years of age, unpublished data). Furthermore, AGM1 and AGM2 previously underwent the same routine sampling procedures one (AGM1: preinfection) and three times (AGM2: preinfection, 7 DPI, and 14 DPI) without complication. In our previous experience fatal complications have only occurred in animals that were severely debilitated at the time of the procedure (one was CD8-depleted-SIV infected and the second was infected with *Mycobacterium tuberculosis* CDC1551). The postmortem findings in these debilitated animals that developed fatal complications were also distinct (no evidence of diffuse alveolar disease) from the two AGMs reported herein. Therefore in our experience, routine sampling procedures do not cause the severe COVID-19 phenotype observed in AGM1 and AGM2, even in the rare cases where fatal complications occur.

Further, we found dramatic increases in plasma cytokines with progression of COVID-19 in NHPs, which further contributes to the development of the COVID-19 phenotypes seen in these infected AGMs and RM. The pathogenesis of ARDS is still poorly understood. ARDS has multiple causes and several animal models have been utilized in the past to study this syndrome. These include models of sepsis, hyperoxia, aerosolized toxin (36), and acid aspiration (37). These models have highlighted the importance of the innate host immune response in the development of acute lung injury. Proinflammatory cytokines including TNFα, IL-1β, IL-8, IL-6, G-CSF, MCP-1, and MIP-1 have been shown to be elevated during the acute phases of acute lung injury (ALI) (38). In human COVID-19, circulating IL-6 has been shown to correlate with radiographic abnormalities of pneumonia (3). Indeed, overexpression of several of these cytokines were observed in both animals that progressed to ARDS. This differed from the cytokine profile in the AGMs and RMs that reached study endpoint. Only a few cytokines (IL-6, IL-8 and IL-10) were elevated in AGM3; whereas, AGM4 exhibited an intermediate phenotype with increased levels of several cytokines (IFNγ, IL-8, IL-13, and IL-4) but only a mild increase in IL-6. Interestingly, at 7 DPI all four AGMs had increased levels of IFNγ, with the two AGMs that progressed having the highest plasma concentration. The IFNγ plasma levels in the AGMs at 7 DPI were positively associated with viral load in bronchial brush samples at the same time (p=0.015, R=0.97, Pearson test) suggesting that viral load may be driving the IFNγ response. Some groups have proposed that IFNγ production may be favorable to the virus through upregulation of ACE2 from IFNγ stimulation (39). Thus, elevated IFNγ in plasma could be explored as a potential predictive biomarker for advanced disease in people.

Several predisposing conditions are known to increase the likelihood of developing severe disease in people following infection with SARS-CoV-2. Age (2), weight (40), and sex (2,41) have been identified as potential predisposing factors for developing severe disease. All of the AGMs included in our study were aged, with an estimated age of 16 years old. Both animals that progressed to severe disease were also female and low weight. This differs from what is reported in COVID-19 patients in which male gender (24,42) and obesity (40) have been shown to have a higher prevalence of severe disease. The AGMs used in this infection study were also imported from nondomestic sources. Detailed longitudinal information (*e.g*. medical history, housing, diet) was not available for the AGMs as it was for the RM that were born at TNPRC. All of the AGMs used in the study were found to be in excellent health upon importation, underwent an unremarkable 90-day quarantine period, and were housed at the TNPRC for 10 months prior to use. All animals were examined, and screened for viral, bacterial, and parasitic infections prior to inclusion as subjects on this study. Although the animals were deemed clinically healthy at the time of initiation of the study, there may have been historical factors that predisposed them to enhanced COVID-19 disease.

Our findings in AGMs differ from previous reports through the identification of a severe phenotype that exhibited rapid clinical decline, acute respiratory distress, and cytokine release concurrently. Major differences in study design may account for the discrepancy between our results and those of prior studies utilizing AGMs including the age of the animals and the strain of virus that was used. Apart from the severe phenotype observed in two of the animals, our findings are otherwise consistent with prior studies with the surviving AGMs showing mild clinical disease, pathology, and prolonged viral shedding (15,18). The RMs in our study also exhibited mild clinical disease and pathology with shorter viral shedding from mucosal sites compared to the AGMs. We did not observe the moderate pathology reported by others, but our study lacked necropsies at early time points wherein the majority of the pulmonary pathology was described. Similarly, we did not observe the cytokine elevations in RM reported by others following SARS-CoV-2 infection (16,17); however, in these prior studies cytokine elevations were acute and transient, and therefore the one week post infection time point used in our study may have been too late to capture these changes. Consistent with previous observations, viral load in both AGMs and RMs was detectable in the lung, mouth, nose, throat, and rectum following infection with SARS-CoV-2 supporting and adding to the prior work of others that show even in the absence of severe disease RMs and AGMs still have utility for testing vaccines and therapeutics that ameliorate disease and blunt viral shedding.

This study demonstrates that following exposure to SARS-CoV-2 aged AGMs develop a spectrum of disease, from mild to severe COVID-19, which in some cases progress to ARDS. The cytokine expression profile in the two animals that developed ARDS is similar to that seen in the severe human disease phenotype. Animal models play a crucial role in elucidating the early pathogenic mechanisms and virus-host interactions for emerging infectious diseases like SARS-CoV-2. Animal models of both mild and severe disease manifestations of COVID-19 are needed to aid with the identification of early clinical and immunological biomarkers that are predictive of mortality and disease severity and to facilitate disease management. Our data suggest that both RM and AGM are capable of modeling mild manifestations of SARS-CoV-2 infection and that aged AGMs may additionally be capable of modeling severe disease manifestations including ARDS. Furthermore, aged AGMs may be very useful for investigating the mechanisms of progression to severe COVID-19 observed with greater frequency in the aging population.

## Methods

### Virus

The virus used for experimental infection was SARS-CoV-2; 2019-nCoV/USA-WA1/2020 (MN985325.1 (43)). Virus stock was prepared in Vero E6 cells and sequence confirmed by deep sequencing. Plaque assays were performed in Vero E6 cells.

### Animals and procedures

A total of eight animals, four aged (≈16 years of age), wild-caught AGM (2M, 2F) and four, adult (13-15 years of age) RM (3M, 1F) were used in this study. Animals (n=4) were exposed to SARS-CoV-2 either by small particle aerosol (44) or multiroute combination. The 4 animals (AGM1, AGM4, RM3, RM4) were exposed by aerosol and received an inhaled dose of approximately 2×10^3^ TCID_50_. The other four animals (AGM2, AGM3, RM1, RM2) were exposed by inoculating a cumulative dose of 3.61×10^6^ PFU through multiple routes (oral, 1 mL; nasal, 1mL; intratracheal, 1 mL; conjunctival, 50 μL per eye). Animals were observed for 21 days including twice daily monitoring. Pre- and postexposure samples included blood, CSF, feces, urine, bronchioalveolar lavage, and mucosal swabs (buccal, nasal, pharyngeal, rectal, vaginal, and bronchial brush). Blood was collected at postexposure days −14, 1, 3 (aerosol) or 4 (multiroute), 7, 14, 21, and at necropsy. CSF, feces, urine, bronchioalveolar lavage, and mucosal swabs were collected at post exposure days −14, 7, 14, 21, and at necropsy. Physical exam, plethysmography, and imaging (radiographs and PET/CT) occurred 7 days prior to exposure and then weekly thereafter. Animals were euthanized for necropsy after three weeks post exposure, or when humane end points were reached.

### Necropsy

Postmortem examination was performed by a board-certified veterinary pathologist. Blood was collected via intracardiac aspiration. Euthanasia was performed by intracardiac installation of 2 mL of sodium pentobarbital. CSF was collected from the atlanto-occipital space (cisterna magna). Mucosal swabs were collected from the oral cavity, nasal cavity, pharynx, rectum, and vagina. The pluck was removed in its entirety. The left and right lungs were photographed and weighed separately. A bronchial brush was used to sample the mainstem bronchi of the right and left lower lobes. Bronchoalveolar lavage was performed on the right caudal lung lobe. Samples from the left anterior and caudal lung lobes were collected fresh and in media for further processing. All right lung lobes were infused and stored in fixative for microscopic evaluation. The remainder of the necropsy was performed routinely with collection of tissues in media, fixative, or fresh.

Tissue samples were fixed in Z-fix (Anatech), embedded in paraffin and 5 um thick sections were cut, adhered to charged glass slides and stained routinely. Tissue examined microscopically included: nasal turbinate, nasopharynx, trachea, carotid artery, aorta, heart, tongue, salivary gland, esophagus, stomach, duodenum, jejunum, pancreas, ileocecal junction, colon (ascending, transverse, descending), rectum, liver, gall bladder, spleen, kidney, urinary bladder, thyroid, pituitary, adrenal, lymph nodes (bronchial, mesenteric, submandibular, cervical, axillary, inguinal, bronchial), tonsils (palatine, lingual), brain (olfactory bulb, frontal cortex, temporal cortex, parietal cortex, occipital cortex, basal ganglia, cerebellum, brainstem), spinal cord (cervical), and reproductive system (ovary, uterus, vagina or testis, seminal vesicle, prostate).

All slides were scanned on a Zeiss Axio Scan.Z1 digital slide scanner. Images and figures were made using HALO software (Indica Labs).

### Histopathologic Scoring

Pulmonary pathology was scored using two separate random forest tissue segmentation algorithms trained by a veterinary pathologist to recognize fibrin and edema and cellular inflammation using HALO software. Tissue sections from each of the right lung lobes was segmented using the trained algorithms to quantify the percentage of tissue effected by fibrin and edema or cellular inflammation. The percentage of inflammation was converted to a pathology score based on the scoring system in the table below. The “Histopathology score” was made by summating the fibrin and edema and cellular inflammation scores for each lobe.

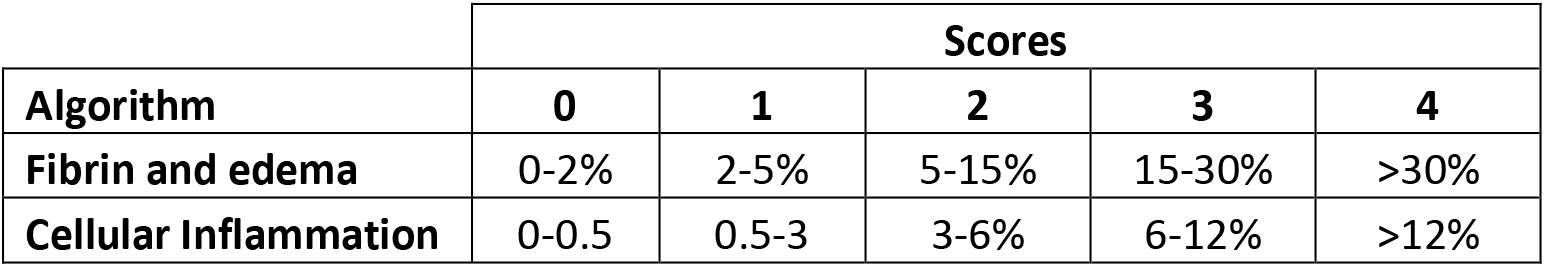

Semiquantitative scores were generated by a veterinary pathologist for lesions within other tissues and specific tissue compartments within the lung. Lesions were scored based on severity as either lacking a lesion (-) or being minimally (+), mildly (++), moderately (+++), or severely (++++) affected.

### Quantification of Swab Viral RNA

Swab and bronchial brush samples were collected in 200 μL of DNA/RNA Shield 1x (Cat.# R1200, Zymo Research, Irvine, CA) and extracted for Viral RNA (vRNA) using the Quick-RNA Viral kit (Cat.# R1034/5, Zymo Research). The Viral RNA Buffer was dispensed directly to the swab in the DNA/RNA Shield. A modification to the manufacturers’ protocol was made to insert the swab directly into the spin column to centrifugate allowing all the solution to cross the spin column membrane. The vRNA was the eluted (45 μL) from which 5 μL was added in a 0.1 mL fast 96-well optical microtiter plate format (Cat #4346906, Thermo Fisher, CA) for a 20 μL RT-qPCR reaction. The RT-qPCR reaction used TaqPath 1-Step Multiplex Master Mix (Cat.# A28527, Thermo Fisher) along with 2019-nCoV RUO Kit (Cat.# 10006713, IDTDNA, Coralville, IA) a premix of forward and reverse primers and a FAM labeled probe targeting the N1 amplicon of N gene of SARS2-nCoV19 (accession MN908947). The reaction master mix were added using an X-stream repeating pipette (Eppendorf, Hauppauge, NY) to the microtiter plates which were covered with optical film (cat. #4311971; Thermo Fisher), vortexed, and pulse centrifuged. The RT-qPCR reaction was subjected to RT-qPCR a program of, UNG incubation at 25°C for 2 minutes, RT incubation at 50°C for 15 minutes, and an enzyme activation at 95°C for 2 minutes followed by 40 cycles of a denaturing step at 95°C for 3 seconds and annealing at 60°C for 30 seconds. Fluorescence signals were detected with an Applied Biosystems QuantStudio 6 Sequence Detector. Data were captured and analyzed with Sequence Detector Software v1.3 (Applied Biosystems, Foster City, CA). Viral copy numbers were calculated by plotting Cq values obtained from unknown (i.e. test) samples against a standard curve representing known viral copy numbers. The limit of detection of the viral RNA assay was 10 copies per reaction volume. A 2019-nCoV positive control (Cat.# 10006625, IDTDNA) were analyzed in parallel with every set of test samples to verify that the RT-qPCR master mix and reagents were prepared correctly to produce amplification of the target nucleic acid. A non-template control (NTC) was included in the qPCR to ensure that there was no cross-contamination between reactions.

### Immunohistochemistry

5um sections of Formalin-fixed, paraffin-embedded lung were mounted on charged glass slides, baked overnight at 56°C and passed through Xylene, graded ethanol, and double distilled water to remove paraffin and rehydrate tissue sections. A microwave was used for heat induced epitope retrieval. Slides were heated in a high pH solution (Vector Labs H-3301), rinsed in hot water and transferred to a heated low pH solution (Vector Labs H-3300) where they were allowed to cool to room temperature. Sections were washed in a solution of phosphate-buffered saline and fish gelatin (PBS-FSG) and transferred to a humidified chamber. Tissues were blocked with 10% normal goat serum (NGS) for 40 minutes, followed by a 60-minute incubation with the primary antibodies (SARS-CoV-2 nucleoprotein, mouse IgG1 (Sino Biological, cat#40143-MM08); ACE2, rabbit polyclonal (Millipore, cat# HPA000288); Iba-1, rabbit polyclonal (Wako, cat# 019-19741); or pancytokeratin, rabbit polyclonal (Dako, cat#Z0622)) diluted in NGS at a concentration of 1:200 and 1:100, respectively). Slides were washed twice in PBS-FSG with Tritonx100, followed by a third wash in PBS-FSG. Slides were transferred to the humidified chamber and incubated, for 40 minutes, with secondary antibodies tagged with Alexa Fluor fluorochromes and diluted 1:1000 in NGS. Following washes, DAPI (4’,6-diamidino-2-phenylindole) was used to label the nuclei of each section. Slides were mounted using a homemade anti-quenching mounting media containing Mowiol (Calbiochem #475904) and DABCO (Sigma #D2522) and imaged with a Zeiss Axio Slide Scanner.

### Cytokine Production in Plasma

Plasma was collected by spinning and was thawed before use. Cytokines were measured using Mesoscale Discovery using a V-Plex Proinflammatory Panel 1, 10-Plex (IFN-γ, IL-1β, IL-2, IL-4, IL-6, IL-8, IL-10, IL-12p70, IL-13, TNF-α) (#K15049D, Mesoscale Discovery, Rockville, Maryland) following the instructions of the kit. The plate was read on a MESO Quick Plex SQ120 machine.

Heatmaps were generated using the ‘pheatmap’ package in R(45,46). Data were normalized by dividing raw values at week 1 and necropsy by baseline values for each animal, followed by the application of log2. Values below the limit of detection were replaced with the lowest limit of detection value based on the standard curve for each run, or with the lowest value detected during the run, whichever was smaller. Polar coordinate plots were generated using the ‘ggplot2’ package in R(47)(46), using the same normalized data shown in the heatmap. Scatterplots were drawn using raw data points and display Pearson’s correlation coefficients.

### Detection of binding IgG antibody in plasma

Serum samples collected at preinfection and at necropsy were tested for binding IgG antibodies against SARS-CoV-2 S1/S2 proteins using an ELISA kit from XpressBio (cat# SP864C). The assays were performed per directions of the manufacturer. In brief, the serum was diluted 1:50 in Sample Diluent. One hundred microliters of diluted serum were pipetted into the wells of the ELISA plate. The plate was covered and incubated at 37° C for 45 min. After incubation, the wells were washed 5 times with 1X wash solution. One hundred microliters of Peroxidase Conjugate were pipetted into each test well. The plate was covered and incubated at 37° C for 45 min. After incubation, the wells were washed 5 times with 1X Wash solution. One hundred microliters of ABTS Peroxidase Substrate was pipetted into each test well. The plate was incubated at room temperature for 30 minutes. The absorbance of the colorimetric reaction was read at 405 nm. Samples were considered as positive if the difference between the absorbance on the positive viral antigen well and the absorbance on the negative control antigen well was greater or equal to 0.300.

Samples collected at preinfection and weekly post-infection until necropsy were tested for detection of binding IgG antibodies against SARS-CoV-2 nucleoprotein (NP) by MFIA COVID-Plex from Charles River Laboratories. The assays were performed per directions of the manufacturer. Briefly, 25 μL of 50-fold diluted samples, or control serum, were added to each well containing 25 μL of bead solution. The plate was covered and incubated at room temperature at 650 rpm orbital shaking for 60 minutes. After incubation, the wells were washed 3 times with 200 μL of MFIA assay buffer. Fifty microliters of MFIA assay buffer plus 50 μL of biotinylated anti-immunoglobulin (BAG) working dilution were added to each well. The plate was covered and incubated at room temperature at 650 rpm orbital shaking for 30 minutes. After incubation, the wells were washed twice as described above, and 50 μL of MFIA assay buffer plus 50 μL of streptavidin-R-phycoerythrin (SPE) were added to each well. The plate was covered and incubated at room temperature at 650 rpm orbital shaking for 30 minutes. After incubation, the plate was washed 3 times as described above, and 125 μL of MFIA assay buffer were added to each well. Plates were read on a Bio-Plex^®^ 200 System (Bio-Rad Laboratories, Hercules, CA). MFIA scores were calculated using Bio-Plex Manager™ Software v6.2 (Bio-Rad) as indicated by Charles River Laboratories. Samples were considered as positive if the MFIA score was greater or equal to 3.0.

### Statistics

Statistical tests were performed with Graphpad prism v8.4.3. The Mann-Whitney U test was used to compare viral load between species and route of exposure. Pearson correlation test was used to test correlation between cytokines and viral load.

### Study approval

The Institutional Animal Care and Use Committee of Tulane University reviewed and approved all the procedures for this experiment. The Tulane National Primate Research Center is fully accredited by the AAALAC. All animals were cared for in accordance with the ILAR Guide for the Care and Use of Laboratory Animals 8^th^ Edition. The Tulane University Institutional Biosafety Committee approved the procedures for sample handling, inactivation, and removal from BSL3 containment.

## Data Availability

The raw data supporting the findings and figures has been placed in a public data repository which can be accessed here: https://figshare.com/s/0436bb616239b57dc007 and will be made public prior to publication. Material requests can be made to the Tulane National Primate Research Center. Approved requests for materials will be released after completion of a material transfer agreement.

## Author Contributions

RVB was the lead pathologist, contributed to study design, analyzed data, and wrote the manuscript. MV performed cytokine assay, composed figures, and contributed to writing the manuscript. LADM was the project veterinarian, contributed to study design, and writing of the IACUC and manuscript. CJR conceived and performed aerosol experiments and contributed to writing the manuscript. KRL contributed to study design, IACUC protocol preparation, clinical examinations, interpretation of clinical data, and manuscript review. MF analyzed cytokine data and made figures. CJM processed and analyzed samples for RT-qPCR, contributed to writing the manuscript. BB collected and analyzed data. KSP, JAP, SCW provided large preparations of deep sequenced virus from the WRCEVA collection. XQ contributed reagents, to the conceptual development of the study, and manuscript writing. CCM designed IHC panels and performed all the staining. GL contributed to study design, provided administrative support, and aided with sample processing and archiving. NG contributed to study design, study coordination, sample processing, and SOP development. BT, TP contributed to sample processing including RT-qPCR, fluids, swabs, and necropsy tissues. CA analysis and interpretation of antibody data and revision of manuscript. MBB analysis and interpretation of antibody data. MP performed antibody testing. PKD processed and analyzed viral load data and contributed to the writing of the manuscript. NJM contributed to study design, analyzed data and contributed to writing the manuscript. AB reviewed and optimized all technical SOPs and was responsible for safety of this study. TF contributed to study design, planning, and writing of the manuscript. RPB contributed to study design, analysis of clinical and imaging results, and writing the manuscript. JR conceived, designed, and supported study, analyzed data, contributed to writing the manuscript.

## Acknowledgements

We would like to acknowledge Natalie Thornburg at the NCIRD for her help acquiring and characterizing the viral stock used in this infection study. We would like to thank the NIH for supporting this work through the TNPRC base grant (P51 OD011104 59) and grant R24 AI120942 to SCW. We would like to thank FastGrant for their funding support to TF.

---

**Supplemental Figure 1.**
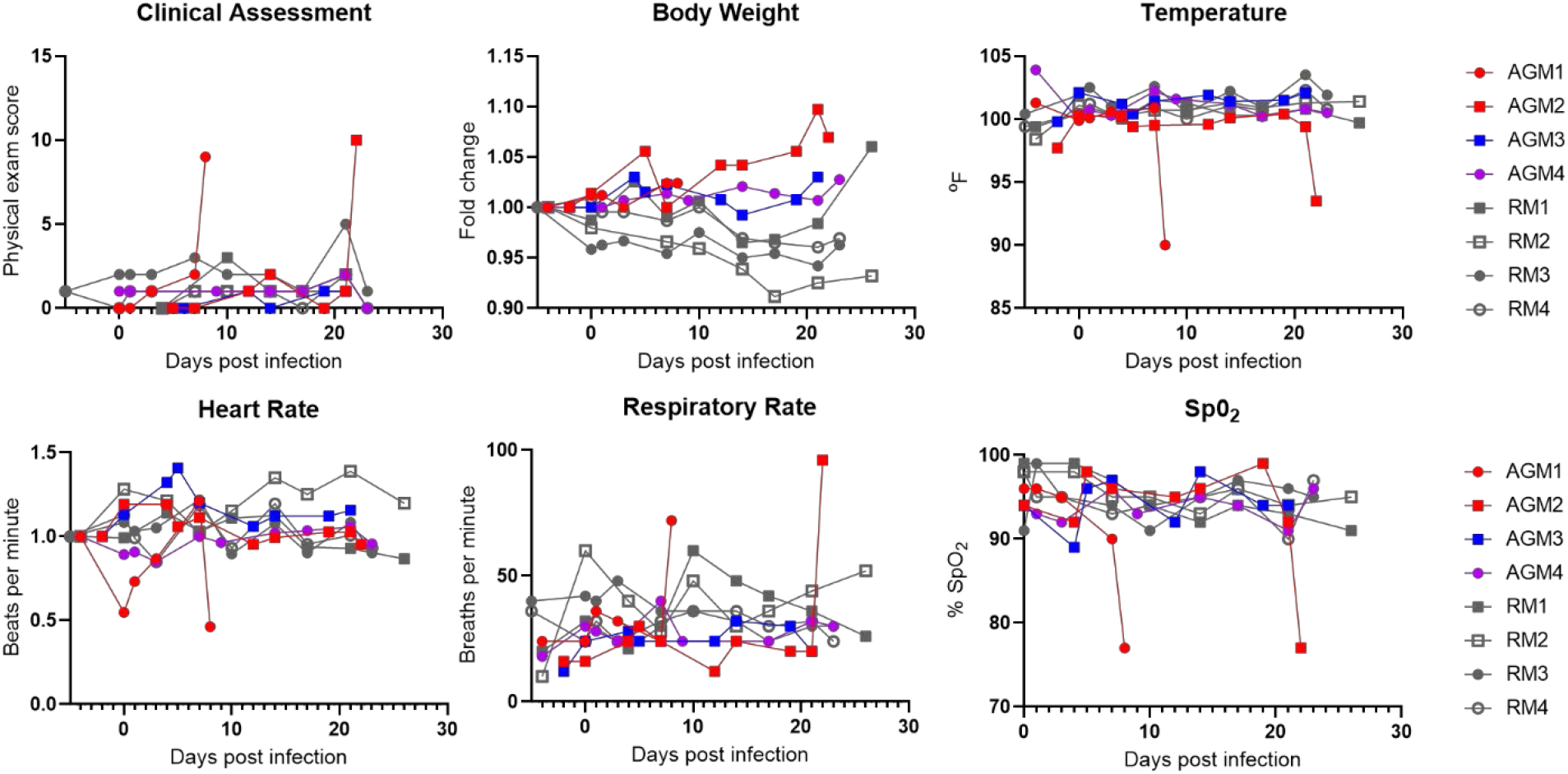
Clinical parameters of African green monkeys and rhesus macaques following exposure to SARS-CoV-2. There were no significant differences in clinical parameters leading up to development of ARDS in the two animals that progressed (red). Progression to ARDS was associated with spike in physical exam scores, respiratory rate, and a dramatic decline in SPO2. Neither weight loss nor fever was associated with SARS-CoV-2 exposure in any of the 8 animals. Circles: aerosol exposure; Squares: multiroute exposure; Gray: Rhesus macaques; Color: AGM by outcome. Red: developed ARDS; Purple: increased cytokines without ARDS; Blue: no cytokine increase or ARDS.

**Supplemental Figure 2.**
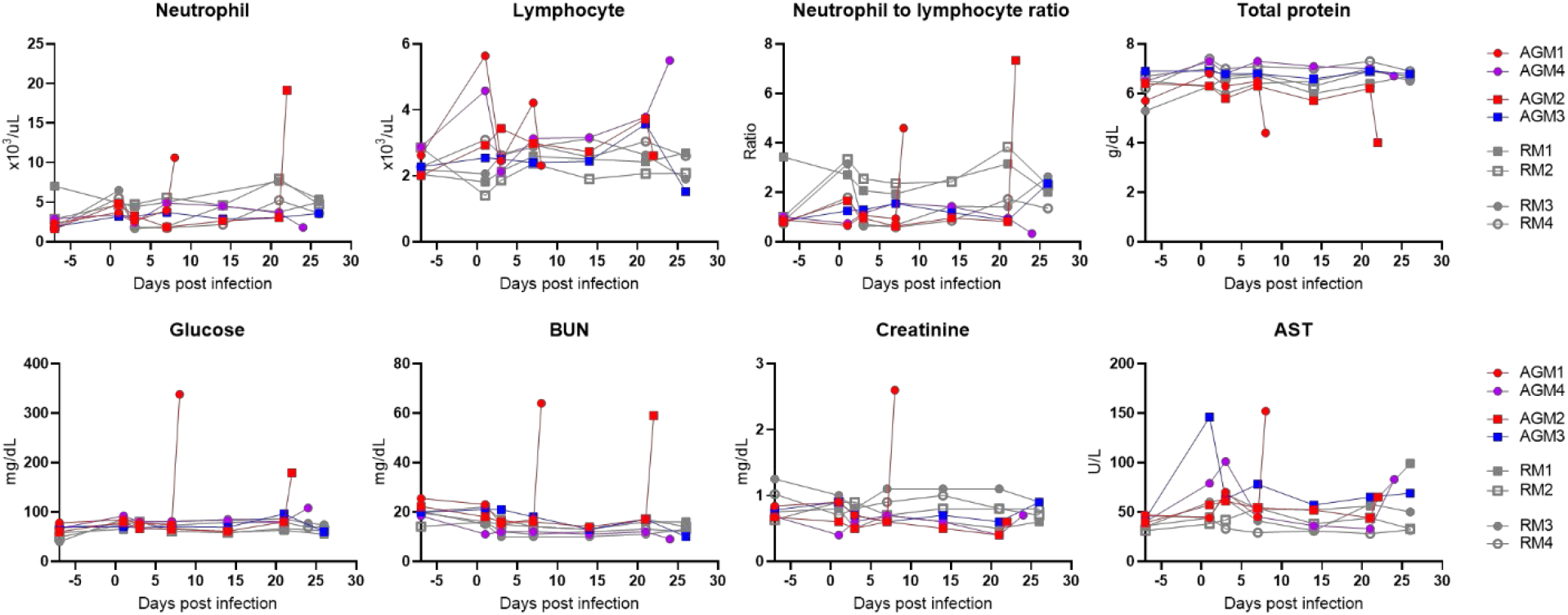
Hematologic and chemistry abnormalities in AGMs with ARDS. The animals that progressed to ARDS (red) only showed hematologic derangements at the terminal timepoint after the onset of respiratory distress. Circles: aerosol exposure; Squares: multiroute exposure; Gray: Rhesus macaques; Color: AGM by outcome. Red: developed ARDS; Purple: increased cytokines without ARDS; Blue: no cytokine increase or ARDS.

**Supplemental Figure 3.**
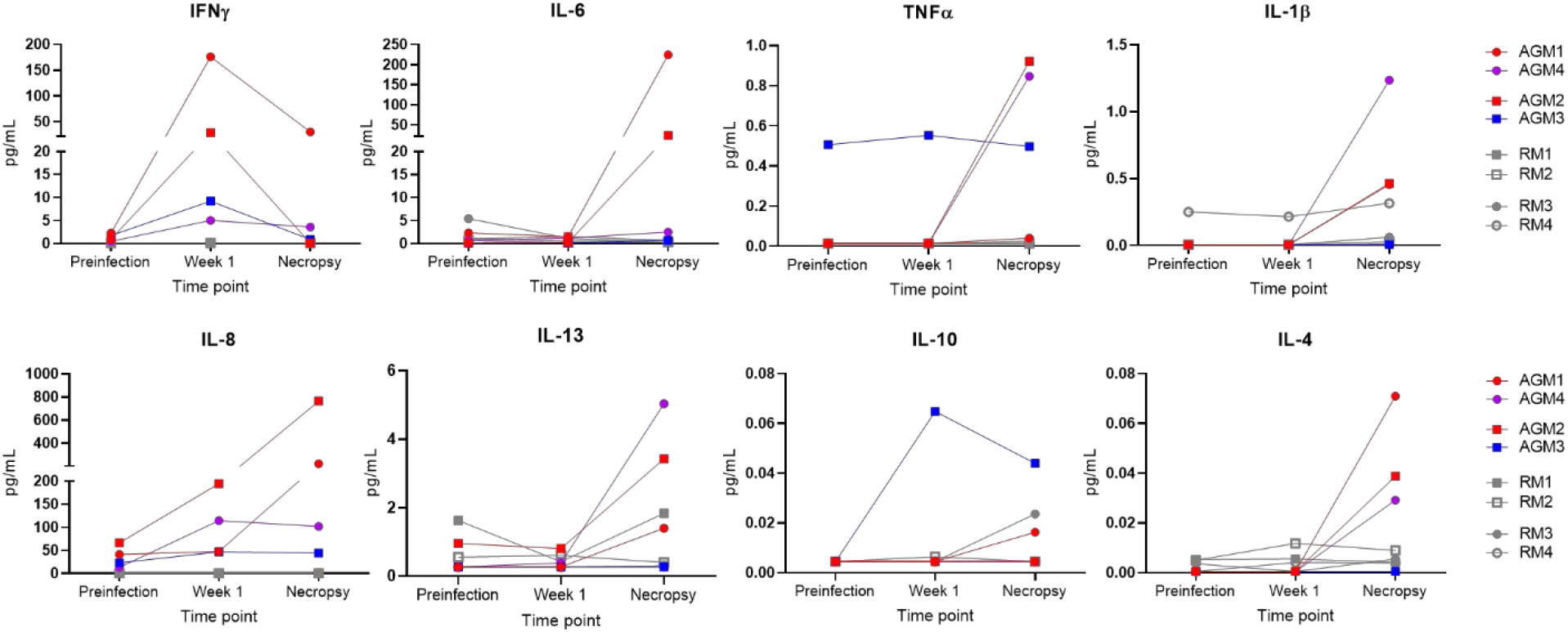
Serum cytokines in SARS-CoV-2 infected African green monkeys and rhesus macaques. AGMs exhibited higher levels of IFNg at one-week post infection compared to rhesus macaques with the two animals (AGM1 and AGM2, red) that progressed showing the highest elevation. At necropsy the two progressors had elevated IL-6, IL-1B, IL-8, IL-13, and IL-4. AGM4 (purple) had a similar cytokine profile except with little elevation in IL-6. AGM3 (blue) had elevated IL-10 one-week post infection and is notable for having the lowest pathology scores of the four AGM. RM showed modest changes in serum cytokines at one-week post infection and at necropsy. Circles: aerosol exposure; Squares: multiroute exposure; Gray: Rhesus macaques; Color: AGM by outcome. Red: developed ARDS; Purple: increased cytokines without ARDS; Blue: no cytokine increase or ARDS.

**Supplemental Figure 4.**
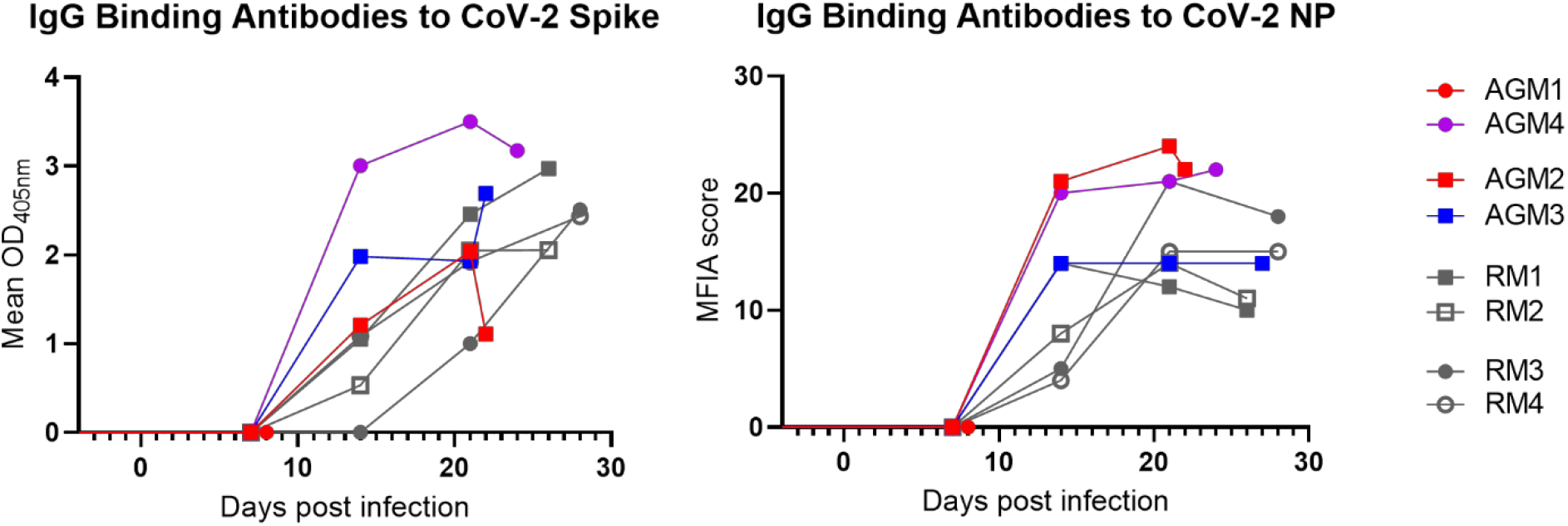
Kinetics of binding antibody responses in SARS-CoV-2 infected African green monkeys and rhesus macaques. Circles: aerosol exposure; Squares: multiroute exposure; Gray: Rhesus macaques; Color: AGM by outcome. Red: developed ARDS; Purple: increased cytokines without ARDS; Blue: no cytokine increase or ARDS.

**Supplemental Figure 5.**
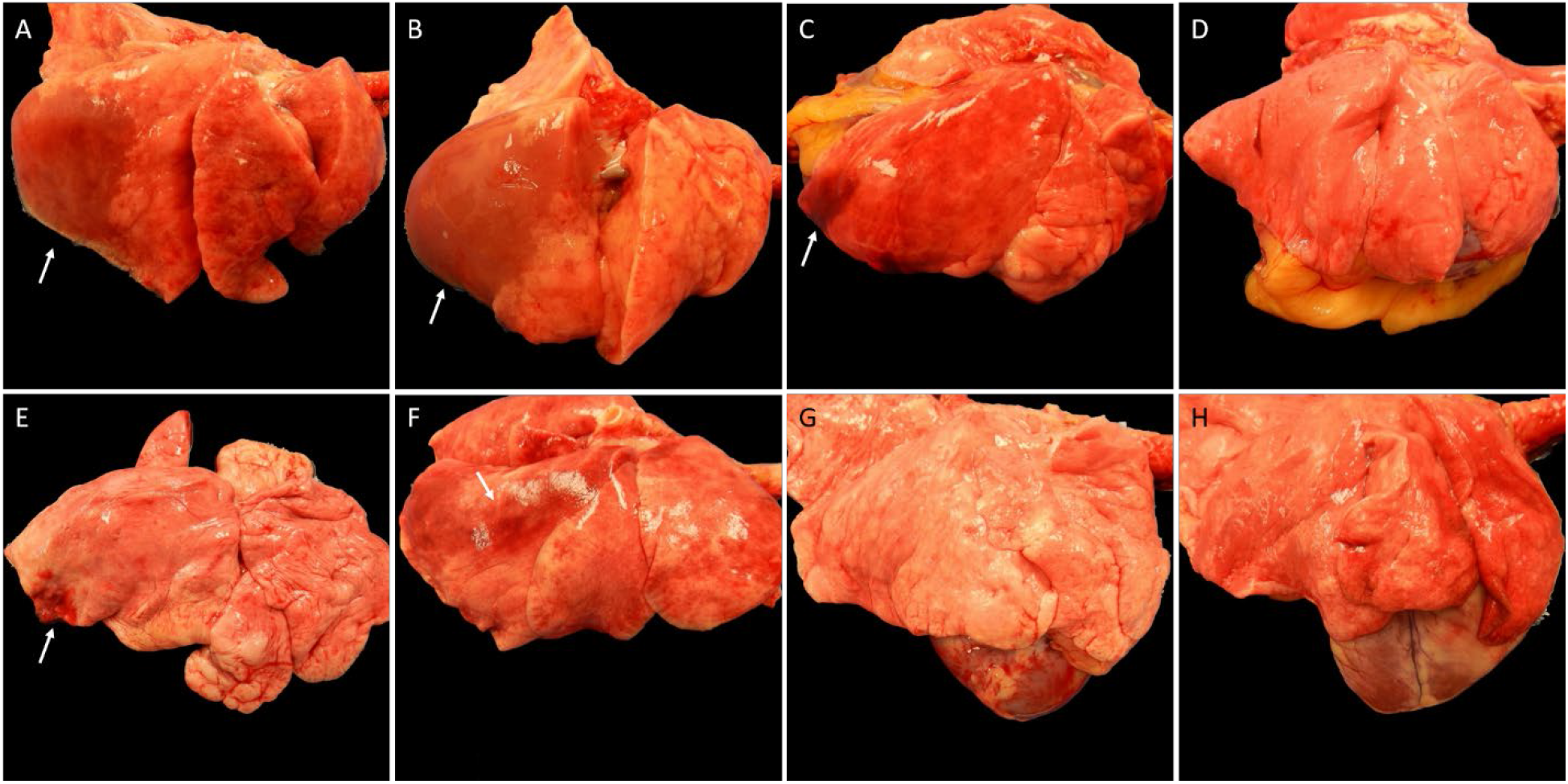
Gross pathology of the right lung lobes of SARS-CoV-2 infected African green monkeys and rhesus macaques. There is extensive consolidation of the right lower lung lobes of AGM1 (**A**) and AGM2 (**B**) with lesser involvement of the middle and anterior lobes. AGM3 (**C**) has focal area of hemorrhage along the caudodorsal margin of the right lower lobe. No gross abnormalities are visible in AGM 4 (**D**). **E-H**, Rhesus macaques. RM1 (**E**) has a scar along the caudodorsal margin of the right lower lobe surrounded by acute hemorrhage. RM2 (**F**) has red mottling of the dorsal margin of the right lower and anterior lung lobes. No gross abnormalities are visible in RM3 and RM4 (**G** and **H**). Arrows point to the lesion described for each animal.

**Supplemental Figure 6.**
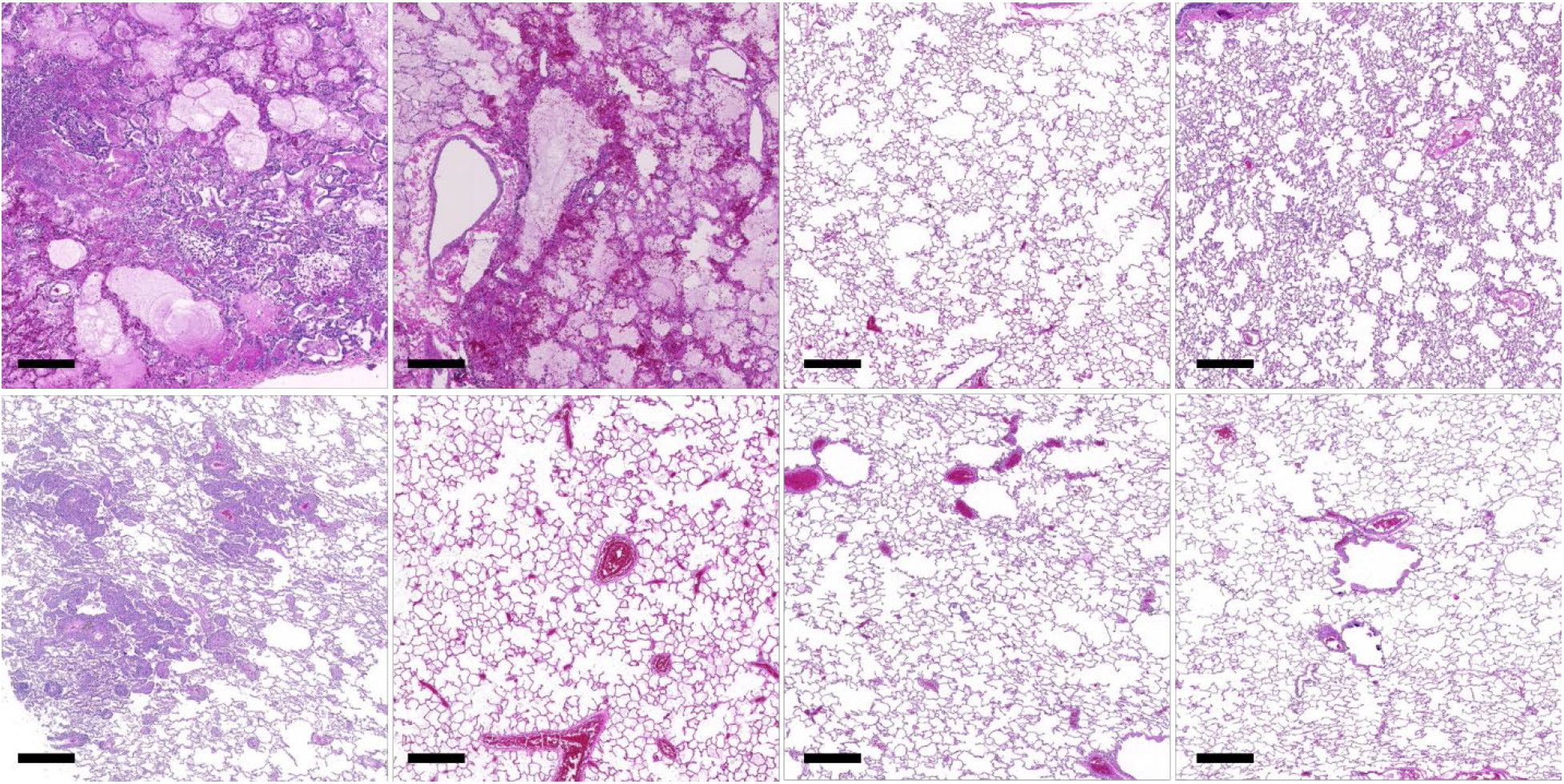
Histopathology of the right caudal lung lobes of SARS-CoV-2 infected African green monkeys and rhesus macaques. Representative images of the histopathology of the right caudal lung lobe from AGM (top row, left to right AGM1-4) and RM (bottom row, left to right RM1-4). H&E, Bar=500um.

**Supplemental Figure 7.**
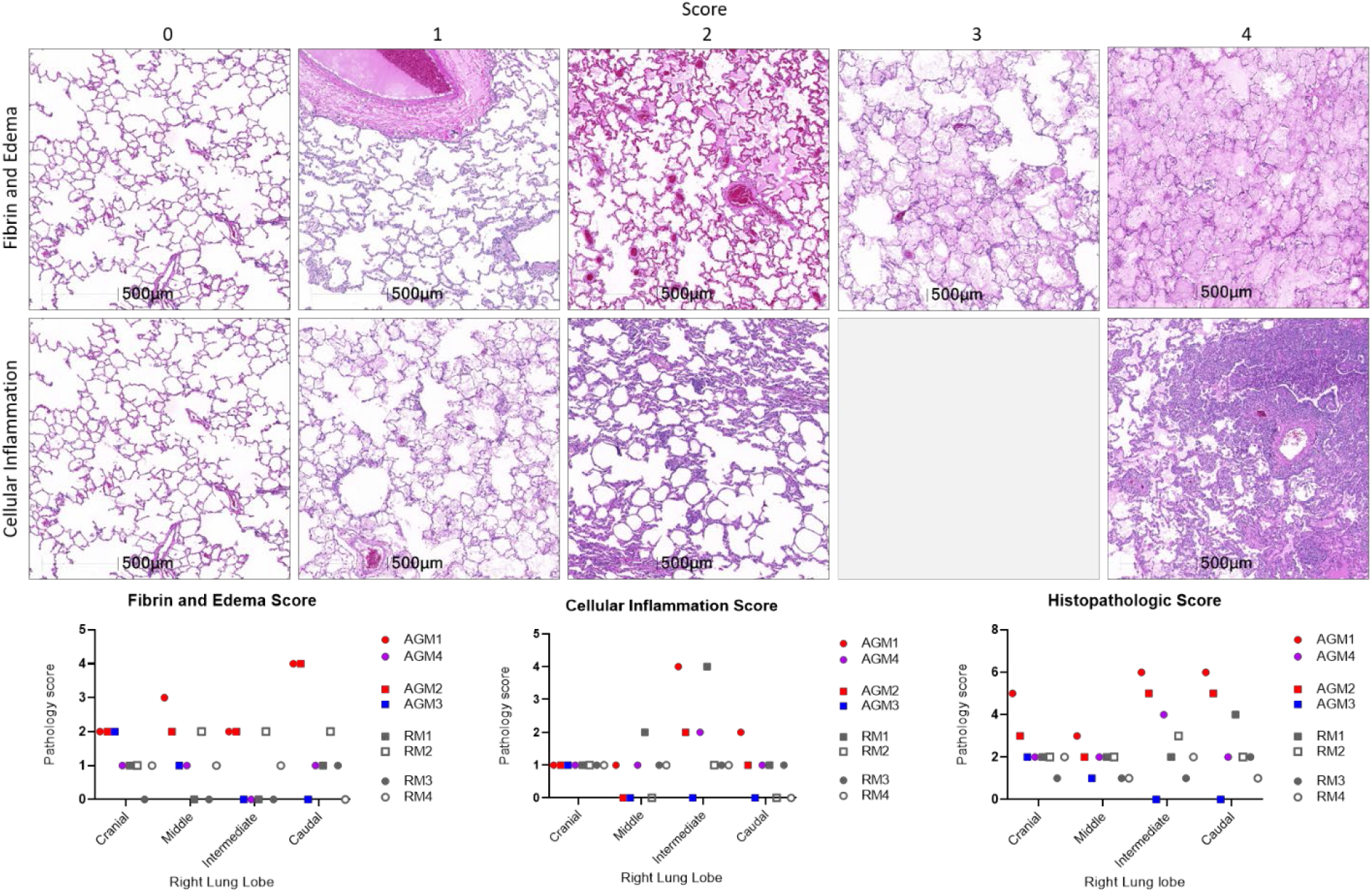
Histopathologic scoring of pulmonary inflammation. Whole slide images of tissue sections from each lung lobe were analyzed for fibrin/edema and cellular inflammation using HALO’s Tissue Classifier module. Top row: representative images of fibrin/edema scores 0-4. Middle row: representative images of cellular inflammation scores 0-2 and 4. Graphs illustrate the score of inflammation in each of the right lung lobes. The Histopathologic Score is the aggregate score of Fibrin/Edema and Cellular Inflammation.

**Supplemental Table 1.** Animal information including species, source, route of exposure, and demographic information from each animal in the study.

| ID | Species | Source | Age (yr) | Sex | Weight (kg) | Exposure (dose) |
| --- | --- | --- | --- | --- | --- | --- |
| AGM1 | *Chlorocebus aethiops sabaesus* | Wild caught, St. Kitts | 16 | F | 4.3 | Aerosol<br>($2 \times 10^3$ TCID<sub>50</sub>) |
| AGM2 | *Chlorocebus aethiops sabaesus* | Wild caught, St. Kitts | 16 | F | 3.9 | Multiroute<br>($3.61 \times 10^6$ PFU) |
| AGM3 | *Chlorocebus aethiops sabaesus* | Wild caught, St. Kitts | 16 | M | 6.9 | Multiroute<br>($3.61 \times 10^6$ PFU) |
| AGM4 | *Chlorocebus aethiops sabaesus* | Wild caught, St. Kitts | 16 | M | 7.5 | Aerosol<br>($2 \times 10^3$ TCID<sub>50</sub>) |
| RM1 | *Macaca mulatta* | Born at TNPRC | 14 | M | 16.7 | Multiroute<br>($3.61 \times 10^6$ PFU) |
| RM2 | *Macaca mulatta* | Born at TNPRC | 13 | F | 6.9 | Multiroute<br>($3.61 \times 10^6$ PFU) |
| RM3 | *Macaca mulatta* | Born at TNPRC | 13 | M | 11.6 | Aerosol<br>($2 \times 10^3$ TCID<sub>50</sub>) |
| RM4 | *Macaca mulatta* | Born at TNPRC | 15 | M | 11 | Aerosol<br>($2 \times 10^3$ TCID<sub>50</sub>) |

**Supplementary Table 2.**
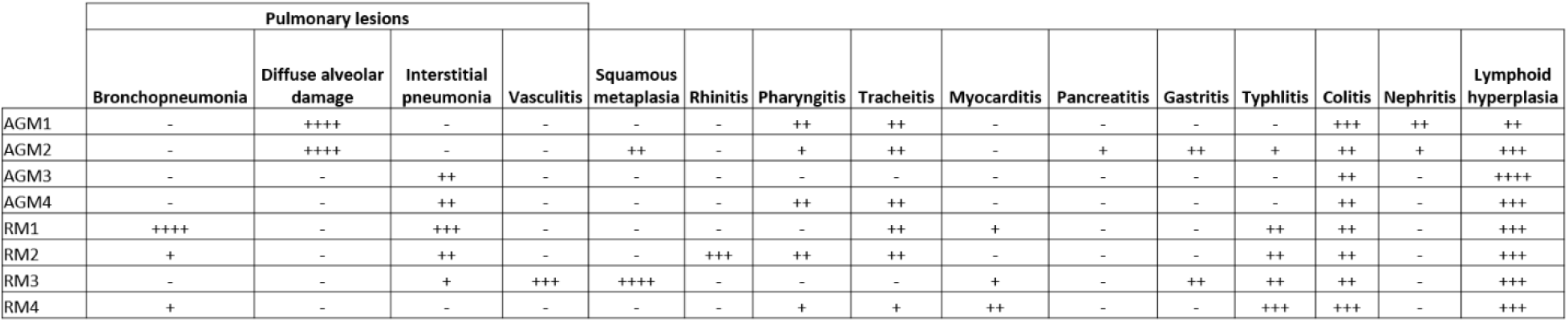
Table of histopathologic findings. Histopathologic changes including localization of inflammation in the lung, morphologic diagnoses, and their semiquantitative severity assigned by a pathologist as follows: – : absent; +: minimal; ++: mild; +++: moderate; ++++: severe. Squamous metaplasia refers to respiratory epithelium of the nasal turbinates.

|  | Pulmonary lesions |  |  |  |  |  |  |  |  |  |  |  |  |  |  |
| --- | --- | --- | --- | --- | --- | --- | --- | --- | --- | --- | --- | --- | --- | --- | --- |
|  | Bronchopneumonia | Diffuse alveolar damage | Interstitial pneumonia | Vasculitis | Squamous metaplasia | Rhinitis | Pharyngitis | Tracheitis | Myocarditis | Pancreatitis | Gastritis | Typhlitis | Colitis | Nephritis | Lymphoid hyperplasia |
| AGM1 | - | ++++ | - | - | - | - | ++ | ++ | - | - | - | - | +++ | ++ | ++ |
| AGM2 | - | ++++ | - | - | ++ | - | + | ++ | - | + | ++ | + | ++ | + | +++ |
| AGM3 | - | - | ++ | - | - | - | - | - | - | - | - | - | ++ | - | ++++ |
| AGM4 | - | - | ++ | - | - | - | ++ | ++ | - | - | - | - | ++ | - | +++ |
| RM1 | ++++ | - | +++ | - | - | - | - | ++ | + | - | - | ++ | ++ | - | +++ |
| RM2 | + | - | ++ | - | - | +++ | ++ | ++ | - | - | - | ++ | ++ | - | +++ |
| RM3 | - | - | + | +++ | ++++ | - | - | - | + | - | ++ | ++ | ++ | - | +++ |
| RM4 | + | - | - | - | - | - | + | + | ++ | - | - | +++ | +++ | - | +++ |

## References

1. Wu Z, McGoogan JM. Characteristics of and Important Lessons From the Coronavirus Disease 2019 (COVID-19) Outbreak in China: Summary of a Report of 72314 Cases From the Chinese Center for Disease Control and Prevention. JAMA. 2020;323(13):1239–1242.

2. Yang X, Yu Y, Xu J, et al. Clinical course and outcomes of critically ill patients with SARS-CoV-2 pneumonia in Wuhan, China: a single-centered, retrospective, observational study. Lancet Respir Med. 2020;8(5):475–481.

3. Chien JY, Hsueh PR, Cheng WC, Yu CJ, Yang PC. Temporal changes in cytokine/chemokine profiles and pulmonary involvement in severe acute respiratory syndrome. Respirology. 2006;11(6):715–722.

4. Channappanavar R, Perlman S. Pathogenic human coronavirus infections: causes and consequences of cytokine storm and immunopathology. Semin Immunopathol. 2017;39(5):529–539.

5. McGonagle D, Sharif K, O’Regan A, Bridgewood C. The Role of Cytokines including Interleukin-6 in COVID-19 induced Pneumonia and Macrophage Activation Syndrome-Like Disease. Autoimmun Rev. 2020;19(6):102537.

6. Weiss SR, Leibowitz JL. Coronavirus pathogenesis. Adv Virus Res. 2011;81:85–164.

7. Lawler JV, Endy TP, Hensley LE, et al. Cynomolgus macaque as an animal model for severe acute respiratory syndrome. PLoS medicine. 2006;3(5):e149.

8. Smits SL, van den Brand JM, de Lang A, et al. Distinct severe acute respiratory syndrome coronavirus-induced acute lung injury pathways in two different nonhuman primate species. Journal of virology. 2011;85(9):4234–4245.

9. Clay CC, Donart N, Fomukong N, et al. Severe acute respiratory syndrome-coronavirus infection in aged nonhuman primates is associated with modulated pulmonary and systemic immune responses. Immun Ageing. 2014;11(1):4.

10. Greenough TC, Carville A, Coderre J, et al. Pneumonitis and multi-organ system disease in common marmosets (Callithrix jacchus) infected with the severe acute respiratory syndrome-associated coronavirus. The American journal of pathology. 2005;167(2):455–463.

11. Datta PK, Liu F, Fischer T, Rappaport J, Qin X. SARS-CoV-2 pandemic and research gaps: Understanding SARS-CoV-2 interaction with the ACE2 receptor and implications for therapy. Theranostics. 2020;10(16):7448–7464.

12. Yeung ML, Yao Y, Jia L, et al. MERS coronavirus induces apoptosis in kidney and lung by upregulating Smad7 and FGF2. Nat Microbiol. 2016;1:16004.

13. Baseler LJ, Falzarano D, Scott DP, et al. An Acute Immune Response to Middle East Respiratory Syndrome Coronavirus Replication Contributes to Viral Pathogenicity. The American journal of pathology. 2016;186(3):630–638.

14. de Wit E, Feldmann F, Cronin J, et al. Prophylactic and therapeutic remdesivir (GS-5734) treatment in the rhesus macaque model of MERS-CoV infection. Proceedings of the National Academy of Sciences of the United States of America. 2020;117(12):6771–6776.

15. Woolsey C, Borisevich V, Prasad AN, et al. Establishment of an African green monkey model for COVID-19. bioRxiv. 2020.

16. Singh DKea. SARS-CoV-2 infection leads to acute infection with dynamic cellular and inflammatory flux in the lung that varies across nonhuman primate species. bioRxiv. 2020.

17. Munster VJ, Feldmann F, Williamson BN, et al. Respiratory disease in rhesus macaques inoculated with SARS-CoV-2. Nature. 2020.

18. Hartman ALea. SARS-CoV-2 infection of African green monkeys results in mild respiratory disease discernible by PET/CT imaging and prolonged shedding of infectious virus from both respiratory and gastrointestinal tracts. bioRxiv. 2020.

19. Rockx B, Kuiken T, Herfst S, et al. Comparative pathogenesis of COVID-19, MERS, and SARS in a nonhuman primate model. Science. 2020;368(6494):1012–1015.

20. van Doremalen N, Lambe T, Spencer A, et al. ChAdOx1 nCoV-19 vaccination prevents SARS-CoV-2 pneumonia in rhesus macaques. bioRxiv. 2020.

21. Gao Q, Bao L, Mao H, et al. Development of an inactivated vaccine candidate for SARS-CoV-2. Science. 2020;369(6499):77–81.

22. Yu J, Tostanoski LH, Peter L, et al. DNA vaccine protection against SARS-CoV-2 in rhesus macaques. Science. 2020.

23. Erasmus JH, Khandhar AP, O’Connor MA, et al. An alphavirus-derived replicon RNA vaccine induces SARS-CoV-2 neutralizing antibody and T cell responses in mice and nonhuman primates. Science translational medicine. 2020.

24. Dudley JP, Lee NT. Disparities in Age-Specific Morbidity and Mortality from SARS-CoV-2 in China and the Republic of Korea. Clinical infectious diseases : an official publication of the Infectious Diseases Society of America. 2020.

25. Zhang X, Cai H, Hu J, et al. Epidemiological, clinical characteristics of cases of SARS-CoV-2 infection with abnormal imaging findings. Int J Infect Dis. 2020;94:81–87.

26. Hu Y, Sun J, Dai Z, et al. Prevalence and severity of corona virus disease 2019 (COVID-19): A systematic review and meta-analysis. J Clin Virol. 2020;127:104371.

27. Wang W, Xu Y, Gao R, et al. Detection of SARS-CoV-2 in Different Types of Clinical Specimens. JAMA. 2020.

28. To KK, Tsang OT, Leung WS, et al. Temporal profiles of viral load in posterior oropharyngeal saliva samples and serum antibody responses during infection by SARS-CoV-2: an observational cohort study. Lancet Infect Dis. 2020;20(5):565–574.

29. Qiu L, Liu X, Xiao M, et al. SARS-CoV-2 is not detectable in the vaginal fluid of women with severe COVID-19 infection. Clinical infectious diseases : an official publication of the Infectious Diseases Society of America. 2020.

30. Shi H, Han X, Jiang N, et al. Radiological findings from 81 patients with COVID-19 pneumonia in Wuhan, China: a descriptive study. Lancet Infect Dis. 2020;20(4):425–434.

31. Liu Y, Du X, Chen J, et al. Neutrophil-to-lymphocyte ratio as an independent risk factor for mortality in hospitalized patients with COVID-19. J Infect. 2020;81(1):e6–e12.

32. Fan BE, Chong VCL, Chan SSW, et al. Hematologic parameters in patients with COVID-19 infection. Am J Hematol. 2020;95(6):E131–E134.

33. Huang C, Wang Y, Li X, et al. Clinical features of patients infected with 2019 novel coronavirus in Wuhan, China. Lancet. 2020;395(10223):497–506.

34. Ferguson ND, Fan E, Camporota L, et al. The Berlin definition of ARDS: an expanded rationale, justification, and supplementary material. Intensive Care Med. 2012;38(10):1573–1582.

35. Raghavendran K, Napolitano LM. Definition of ALI/ARDS. Crit Care Clin. 2011;27(3):429–437.

36. Roy CJ, Ehrbar DJ, Bohorova N, et al. Rescue of rhesus macaques from the lethality of aerosolized ricin toxin. JCI insight. 2019;4(1).

37. Matute-Bello G, Frevert CW, Martin TR. Animal models of acute lung injury. Am J Physiol Lung Cell Mol Physiol. 2008;295(3):L379–399.

38. O’Grady NP, Preas HL, Pugin J, et al. Local inflammatory responses following bronchial endotoxin instillation in humans. Am J Respir Crit Care Med. 2001;163(7):1591–1598.

39. Ziegler CGK, Allon SJ, Nyquist SK, et al. SARS-CoV-2 Receptor ACE2 Is an Interferon-Stimulated Gene in Human Airway Epithelial Cells and Is Detected in Specific Cell Subsets across Tissues. Cell. 2020;181(5):1016–1035 e1019.

40. Simonnet A, Chetboun M, Poissy J, et al. High Prevalence of Obesity in Severe Acute Respiratory Syndrome Coronavirus-2 (SARS-CoV-2) Requiring Invasive Mechanical Ventilation. Obesity (Silver Spring). 2020;28(7):1195–1199.

41. Walter LA, McGregor AJ. Sex- and Gender-specific Observations and Implications for COVID-19. West J Emerg Med. 2020;21(3):507–509.

42. Jin JM, Bai P, He W, et al. Gender Differences in Patients With COVID-19: Focus on Severity and Mortality. Front Public Health. 2020;8:152.

43. Harcourt J, Tamin A, Lu X, et al. Severe Acute Respiratory Syndrome Coronavirus 2 from Patient with Coronavirus Disease, United States. Emerging infectious diseases. 2020;26(6):1266–1273.

44. Hartings JM, Roy CJ. The automated bioaerosol exposure system: preclinical platform development and a respiratory dosimetry application with nonhuman primates. J Pharmacol Toxicol Methods. 2004;49(1):39–55.

45. Kolde R. pheatmap: Pretty Heatmaps. R package version 1012. 2018.

46. Team RC. R: A language and environment for statistical computing. R Foundation for Statistical Computing. 2019.

47. Wickham H. ggplot2: Elegant Graphics for Data Analysis. 2016.

